# iSUMO - integrative prediction of functionally relevant SUMOylation events

**DOI:** 10.1101/056564

**Authors:** Xiaotong Yao, Shuvadeep Maity, Shashank Gandhi, Marcin Imielenski, Christine Vogel

## Abstract

Post-translational modifications by the Small Ubiquitin-like Modifier (SUMO) are essential for diverse cellular functions. Large-scale experiment and sequence-based predictions have identified thousands of SUMOylated proteins. However, the overlap between the datasets is small, suggesting many false positives with low functional relevance. Therefore, we integrated ~800 sequence features and protein characteristics such as cellular function and protein-protein interactions in a machine learning approach to score likely functional SUMOylation events (iSUMO). iSUMO is trained on a total of 24 large-scale datasets, and it predicts 2,291 and 706 SUMO targets in human and yeast, respectively. These estimates are five times higher than what existing sequence-based tools predict at the same 5% false positive rate. Protein-protein and protein-nucleic acid interactions are highly predictive of protein SUMOylation, supporting a role of the modification in protein complex formation. We note the marked prevalence of SUMOylation amongst RNA-binding proteins. We validate iSUMO predictions by experimental or other evidence. iSUMO therefore represents a comprehensive tool to identify high-confidence, functional SUMOylation events for human and yeast.

## Introduction

The covalent attachment of Small Ubiquitin-like Modifier (SUMO) is, based on its common occurrence and wide array of functions in eukaryotic cells, one of the most important post-translational modifications. SUMOylation has been studied from numerous perspectives since its discovery in 1997 ^1^. It is widely conserved across eukaryotes ^2-4^, and in many cases essential for the organismal viability ^5^. SUMOylation resembles ubiquitination in terms of structure, enzymatic pathway, and it has a broad functional spectrum, ranging from chromatin organization ^6^, DNA damage repair ^7^, regulation of transcription ^8^, ribosome biogenesis ^9-10^, messenger RNA (mRNA) processing ^11-12^, nucleus-cytoplasm transport ^13^, to protein localization ^14^, proteolysis (where it cross-talks with ubiquitination)^15^, stress response ^16^ and other functions^17^.

Several computational approaches exist that predict SUMOylation based on the conserved amino acid sequence motif Ψ-K-X-D/E, where Ψ is a hydrophobic residue, K is the lysine being modified, X is any amino acid, and D/E is an acidic residue ^18-21^. However, these sequence-based predictions have many false positives and false negatives: when comparing them to experimental data, the intersection is only small. For example, half of the human proteins contain the above SUMOylation motif in their sequence, but the modification is verified for only a small fraction. In addition, recent experimental data suggests that SUMOylation may also act on motifs other than the one described above ^22^, highlighting the need for methods that move beyond use of sequence alone.

Several experimental methods have been developed to identify SUMO-targets. For example, immunoprecipitation using antibodies against SUMOylated proteins reveals SUMO conjugation with high confidence ^23^, but the assay only works with a small number of proteins at a time. In comparison, mass spectrometry based methods sample a large fraction of the proteome and have by now identified thousands of SUMO targets in yeast and human (**Figure 1**). However, it is often unclear what the false-positive and false-negative identification rates of these approaches are. For example, a recent and in-depth screen of human SUMOylation targets using advanced technology identified only 1,606 proteins ^22^, and overlap between this and other studies is small (**Figure 1**). In comparison, when using prediction tools that are sole based on sequence features, many more targets have been identified, e.g. 9,173 of 16,849 human proteins (54%) are predicted to have a SUMOylation motif in their sequence.

**Figure 1.**
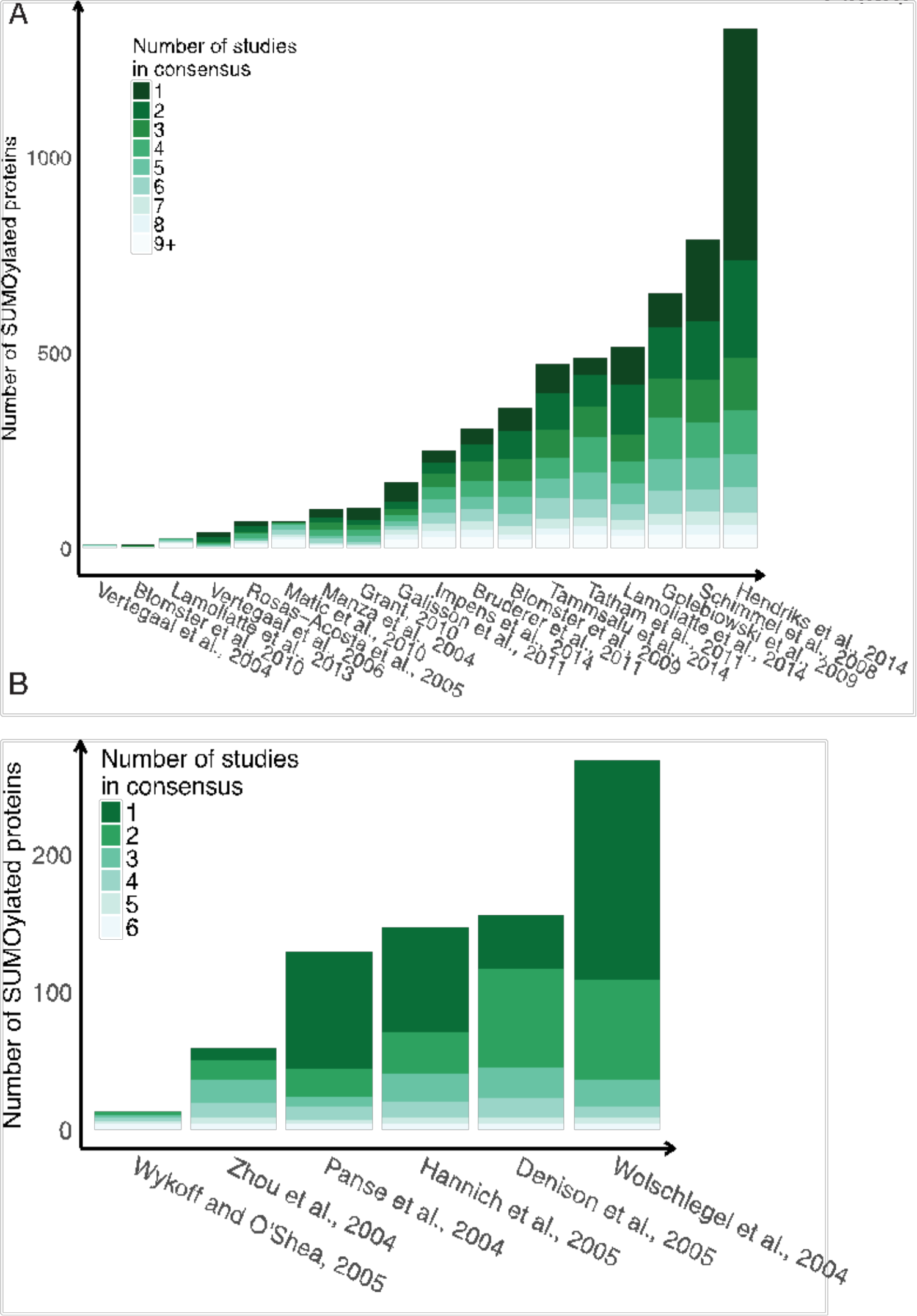
Large-scale experimental datasets for SUMOylated proteins. We assembled 18 and six datasets with mass spectrometry-based identifications of SUMOylated proteins in (A) human and (B) yeast, respectively, which are used as training data in the iSUMO prediction tool. Each column represents a published dataset. Colored entries represent proteins observed as SUMOylated by the respective dataset. Datasets are lists in full in **Suppl. Tables S1**.

Overall, these findings suggest that interpreting large scale identification results remains challenging, as it is likely enriched in many false-positive identifications. To achieve a unified interpretation, we developed iSUMO, a random forest-based predictive model, exploiting the putative SUMOylated proteins discovered in large-scale experiments in their relationship to various protein annotations. iSUMO’s results not only provide a set of SUMOylation events that are likely functional, they also highlight properties common to SUMOylated proteins. In total, these results suggest that ~20% and ~10% of human and yeast proteins are SUMOylated.

## Materials and methods

### Training data sets of experimentally observed SUMOylated proteins

We assembled the results from 18 and 6 large-scale, experimental studies in human ^22, 24-32^ and yeast ^33-38^, respectively, which mapped SUMOylated proteins using mass spectrometry. **Figure 1** summarizes the composition of consensus or unique findings among the studies, and detailed description of the data can be found in **Supplementary Tables S1a and b**. We obtained a total of 2,216 and 555 distinct, well-annotated human and yeast proteins, respectively, to construct a highly quality training dataset.

To integrate the different datasets, we downloaded both reference proteomes from UniProt Reference Proteome database (Homo sapiens, UP000005640; Saccharomyces cerevisiae S288c, UP000002311), and only retained the subset that has been reviewed by Swiss-Prot database. We then filtered both reference proteomes to contain only one protein per gene, restricting unique identity across the combination of Ensembl Gene ID, National Center for Biotechnology Information (NCBI) Gene ID, and UniProt Knowledge Base (UniProtKB)^39^ Accession ID. To avoid introducing biases with dubious function annotations for human genes, we created a high-confidence dataset for the human proteome (>11,000 proteins, **Suppl. Table S6a**). We also present iSUMO predictions with the extended dataset (>16,000 proteins, **Suppl. Table S6b**). For yeast, we used ~6,700 proteins. Within these reference proteomes, we labeled the proteins as “SUMOylated” if they are discovered as SUMOylated in any of the 24 studies described above.

### Sequence-based prediction of protein SUMOylation

For genome-wide prediction of SUMOylation based on protein sequence, we used the Group-based Prediction System-SUMO (GPS-SUMO) _^19, 40^ to predict consensus and non-consensus SUMOylation motifs and SUMO-interaction motifs (SIM) with the threshold set to ‘high’.

### Function enrichment analysis

GO enrichment analysis was carried out using gProfileR package ^41-42^. The query lists were defined as the union of all lists of observed SUMOylated proteins and the background lists are the respective reference proteomes. We set additional parameters to: 1) excluding the GO associations based only on electronic annotation; 2) using adjusted p-values that correct for multiple hypotheses testing; 3) non-ordered query list, thus adopting hypergeometric test. Next we only retained the significant enriched GO terms (false discovery rate<0.05), whose total size are less than 2,000 for human and size within 5 to 1,000 for yeast. Finally, we strictly excluded any term containing “sumo” in its name, case-insensitively, from subsequent modelling steps.

### Attribute selection for predictive modeling

The attributes used for the modeling were taken from multiple sources. The full list of attributes used for predictive modeling is in **Supplementary Table S2**. The attributes include, for example, numbers of occurrences of sequence-based predictions of SUMOylation or SUMO-interaction motifs. These attributes are purely based on sequence predictions and were also used as baseline to compare our own iSUMO predictions. iSUMO also includes attributes derived from associations with Gene Ontology (GO) terms (**Table 1**). Besides, we also incorporated protein-protein interaction data from the STRING database using only the “binding” type activity. We simplified its rich structure into the degree of proteins within this network ^43-45^. Since the taxonomy id 559292 is absent in STRING, we used the equivalent data of 4932 for yeast. Moreover, using the annotations from CORUM database of mammalian protein complex and Costanzo et al., 2016 for yeast ^46^, we counted the number of protein complexes to which each protein belongs and the average size of these complexes (size defined as the number of distinct protein species per complex). Finally, we included other PTM sites of a protein based on the annotations in PhosphoSitePlus for human and dbPTM for yeast ^47-48^, both experimentally supported only, summarizing the number of amino acids that are phosphorylated, acetylated, methylated, or ubiquitylated.

**Table 1.** SUMOylated proteins are biased in their functions. We tested for function enrichment using hypergeometric tests of Gene Ontology (GO) term in human and yeast, respectively. ‘Domain’ codes indicate the three main branches of Gene Ontology: biological processes (BP), cellular compartment (CC), and molecular function (MF). ‘Term size’ refers to the total number of genes associated with the term in the GO database, and ‘Intersection size’ is the intersection with SUMOylated proteins. The ‘Adjusted p-value’ has been corrected for multiple hypotheses testing. The entries are sorted according to the adjusted p-value. Extended data is in **Suppl. Table S2.**

| *A. Homo sapiens* |  |  |  |  |
| --- | --- | --- | --- | --- |
| Domain | Term name | Term size | Intersect ion size | Adjusted P-value |
| BP | RNA processing | 620 | 380 | 9.17E-119 |
|  | mRNA metabolic process | 479 | 293 | 2.87E-89 |
|  | ribonucleoprotein complex biogenesis | 311 | 213 | 5.90E-77 |
|  | ribosome biogenesis | 202 | 149 | 1.18E-59 |
|  | rRNA processing | 170 | 134 | 1.57E-59 |
|  | RNA splicing | 294 | 187 | 4.30E-59 |
|  | mRNA splicing, via spliceosome | 238 | 162 | 5.51E-57 |
|  | ncRNA metabolic process | 382 | 216 | 8.65E-56 |
|  | chromatin organization | 457 | 230 | 1.74E-47 |
|  | regulation of transcription, DNA-templated | 1849 | 610 | 5.25E-47 |
|  | SRP-dependent cotranslational protein targeting to membrane | 83 | 77 | 4.40E-44 |
| CC | chromosome | 604 | 337 | 4.22E-88 |
|  | ribonucleoprotein complex | 470 | 288 | 6.77E-88 |
|  | nuclear chromosome | 388 | 227 | 1.14E-62 |
|  | nucleolus | 558 | 282 | 7.94E-60 |
|  | nucleoplasm part | 491 | 258 | 1.02E-58 |
|  | chromatin | 310 | 180 | 3.46E-48 |
| MF | nucleic acid binding | 1949 | 1014 | 4.45E-281 |
|  | poly(A) RNA binding | 906 | 596 | 3.42E-220 |
|  | RNA binding | 1093 | 665 | 1.07E-219 |
|  | DNA binding | 963 | 428 | 3.95E-72 |
| *B. Saccharomyces cerevisiae* |  |  |  |  |
| Domain | Term name | Term size | Intersect ion size | Adjusted P-value |
| BP | transcription, DNA-templated | 678 | 163 | 3.98E-34 |
|  | organic cyclic compound biosynthetic process | 918 | 194 | 1.64E-33 |
|  | nucleobase-containing compound biosynthetic process | 823 | 181 | 4.25E-33 |
|  | chromatin organization | 325 | 106 | 9.56E-33 |
|  | heterocycle biosynthetic process | 883 | 188 | 1.24E-32 |
|  | macromolecular complex subunit organization | 867 | 173 | 1.6E-25 |
| CC | nuclear lumen | 764 | 168 | 4.58E-30 |
|  | nucleoplasm | 272 | 78 | 4.41E-19 |
|  | chromatin | 141 | 52 | 2.74E-17 |
|  | SWI/SNF superfamily-type complex | 59 | 33 | 4.82E-17 |
| MF | nucleic acid binding | 879 | 160 | 1.81E-18 |
|  | DNA binding | 376 | 75 | 7.31E-09 |
|  | macromolecular complex binding | 203 | 47 | 5.42E-07 |
|  | RNA binding | 524 | 88 | 1.43E-06 |

### Fitting, evaluating, and applying random forest model

iSUMO models SUMOylation state of proteins as a binary classification task based on training data, which comprises categorical and numerical attributes described above. We used the H2O open source machine learning platform with its R interface. The core classification method is random forest. We split the full dataset into 65% training, 20% validation, and 15% testing. We attempted different settings for the training, such as hard setting number of trees versus dynamic termination rule, and restricting different max numbers of tree depth. The model selection is based on the performances with validation set and 10-fold cross validation, considering metrics including area under the receiver operating curve (AUROC), and maximum F1 and F2 values. As training labels were strongly biased towards non-SUMOylated proteins, we also tried balancing the training examples by over sampling of the minority label. Plus, H2O measures relative importance of variables in random forest models by calculating how often a variable is selected at a split of a tree and the amount of improvement in squared error over all trees (http://www.h2o.ai).

## Results

### Many large-scale studies of the SUMOylated proteome exist, but their overlap is small

To obtain a comprehensive training dataset of true-positive SUMOylated proteins, we integrated 18 and six large-scale, experimental studies for human and yeast, respectively. These studies identified individually between ten to >1,600 SUMOylated human proteins, and 13 to 271 SUMOylated yeast proteins. When mapped to the selected reference proteome, we retained a total of 2,270 and 527 SUMO targets for the two species, respectively. About half (45%) of these proteins were identified by two or more studies. In yeast, only one third (30%) of the 527 total SUMOylated proteins were identified by more than one study (**Figure 1**). The lack of overlap between individual studies indicates that the individual studies many have false-positive identifications which are not biologically functional.

### SUMOylated human and yeast proteins bind and process nucleic acids

**Table 1** shows representative, highly significant Gene Ontology (GO) enrichments for both the human and yeast SUMOylated proteins at a (false discovery rate <e-40). The complete results are in the **Suppl. Table S2**. Functions related to DNA processing and metabolism were enriched in both human and yeast SUMOylated proteins, including chromatin organization, DNA damage response, mitosis cell cycle, and transcription; cellular compartments including nucleus, nucleoplasm, nuclear body, and nucleolus. These functions are consistent with our current understanding of SUMOylation’s important role in gene expression regulation ^49-50^. More than two thirds of the 174 human involved in viral gene expression are SUMOylated, which is consistent with the process in which viruses take advantage of host cell SUMOylation to optimize viral gene expression ^51^.

SUMOylated proteins were enriched in functions related to RNA processing, often more significantly than those concerning DNA processing (**Table 1**). The effect was stronger in human than in yeast. The enriched functions included RNA processing, translation elongation and termination, cellular compartments like nucleolus, ribonucleoprotein complex, spliceosomal complex, cytoplasmic ribosomes, and molecular functions like RNA binding – all related to various steps of RNA synthesis and processing.

### Large protein-RNA complexes are extensively SUMOylated

One distinct pattern of SUMOylation was the simultaneous modification of several subunits, probably assisting with organized recruitment of complex components, for instance in proteins in DNA double strand break repair by homologous recombination ^52^. Thus, we first tested if SUMOylated proteins are more often part of any stable protein complex than non-SUMOylated complexes. Indeed, this is the case in both human and yeast (unadjusted hypergeometric test p-value at 1E-132 and 9E-58, respectively).

Next, we tested as if the complexes were enriched in specific functions (**Table 2**). Since most complexes have few subunits, we limited this analysis to the complexes with size larger than 20. The top SUMO-enriched complexes function in, for example, the pre-rRNA complex, involved in ribosome biogenesis, splicing, translation, protein degradation (proteasome), and centromere chromatin complex. We also observed cases whose SUMOylation has so far received little attention. For example, three-quarters (86/110) of the human proteins annotated as part of the signal recognition particle and its co-translational protein targeting to membranes are SUMOylated – which has, to the best of our knowledge, not yet been reported in literature.

**Table 2.** Large protein complexes are heavily SUMOylated. The table shows the subunit compositions of stable protein complexes in human. The upper and lower tables show the largest complexes that are SUMOylated and not SUMOylated, respectively, as determined by the adjusted p-value. The complexes are sorted by descending total number of subunits. The adjusted p-value reports the bias with respect to SUMOylation of the complex subunits. Even complexes with only few RNA-binding subunits, like PA700-20S-PA28 can be SUMOylated.

| Complex name | Complex size | Number of RNA-binding subunits | Number of SUMOylated subunits | Adjusted P-value, SUMO enrichment |
| --- | --- | --- | --- | --- |
| **SUMOylated complexes** |  |  |  |  |
| Spliceosome | 143 | 116 | 106 | 4.37E-29 |
| Nop56p-associated pre-rRNA complex | 104 | 96 | 95 | 1.46E-41 |
| Ribosome, cytoplasmic | 81 | 77 | 71 | 4.83E-28 |
| C complex spliceosome | 80 | 69 | 65 | 1.52E-21 |
| 60S ribosomal subunit, cytoplasmic | 47 | 46 | 42 | 4.44E-17 |
| CEN complex | 37 | 9 | 25 | 1.09E-04 |
| PA700-20S-PA28 complex | 36 | 4 | 28 | 7.12E-08 |
| 17S U2 snRNP | 33 | 30 | 29 | 4.27E-11 |
| 40S ribosomal subunit, cytoplasmic | 31 | 30 | 28 | 1.82E-11 |
| CDC5L complex | 30 | 23 | 23 | 4.68E-06 |
| **Non-SUMOylated complexes** |  |  |  |  |
| 55S ribosome, mitochondrial | 78 | 46 | 1 | 1.00E+00 |
| 39S ribosomal subunit, mitochondrial | 48 | 28 | 0 | 1.00E+00 |
| Mediator complex | 32 | 0 | 6 | 1.00E+00 |
| 28S ribosomal subunit, mitochondrial | 30 | 18 | 1 | 1.00E+00 |
| MLL1-WDR5 complex | 27 | 4 | 15 | 7.86E-01 |
| BRCA1-RNA polymerase II complex | 26 | 9 | 9 | 1.00E+00 |
| TNF-alpha/NF-kappa B signaling complex<br>5 | 25 | 4 | 10 | 1.00E+00 |
| RNA polymerase II holoenzyme complex | 24 | 8 | 8 | 1.00E+00 |
| Emerin complex 32 | 22 | 2 | 14 | 1.25E-01 |
| RNA polymerase II complex | 19 | 2 | 6 | 1.00E+00 |

For both organisms, we found the sizes of complexes tend to be significantly larger when the complex contains both RNA binding and SUMOylated proteins (**Figure 2A**). We then plot the number of SUMOylated subunits against the number of RNA binding proteins per complex, with the size of points corresponding to the size of the complexes (**Figure 2B**). While there are some outliers, there is a clear positive correlation between number of RNA binding and SUMOylated proteins among the largest complexes for both organisms.

**Figure 2.**
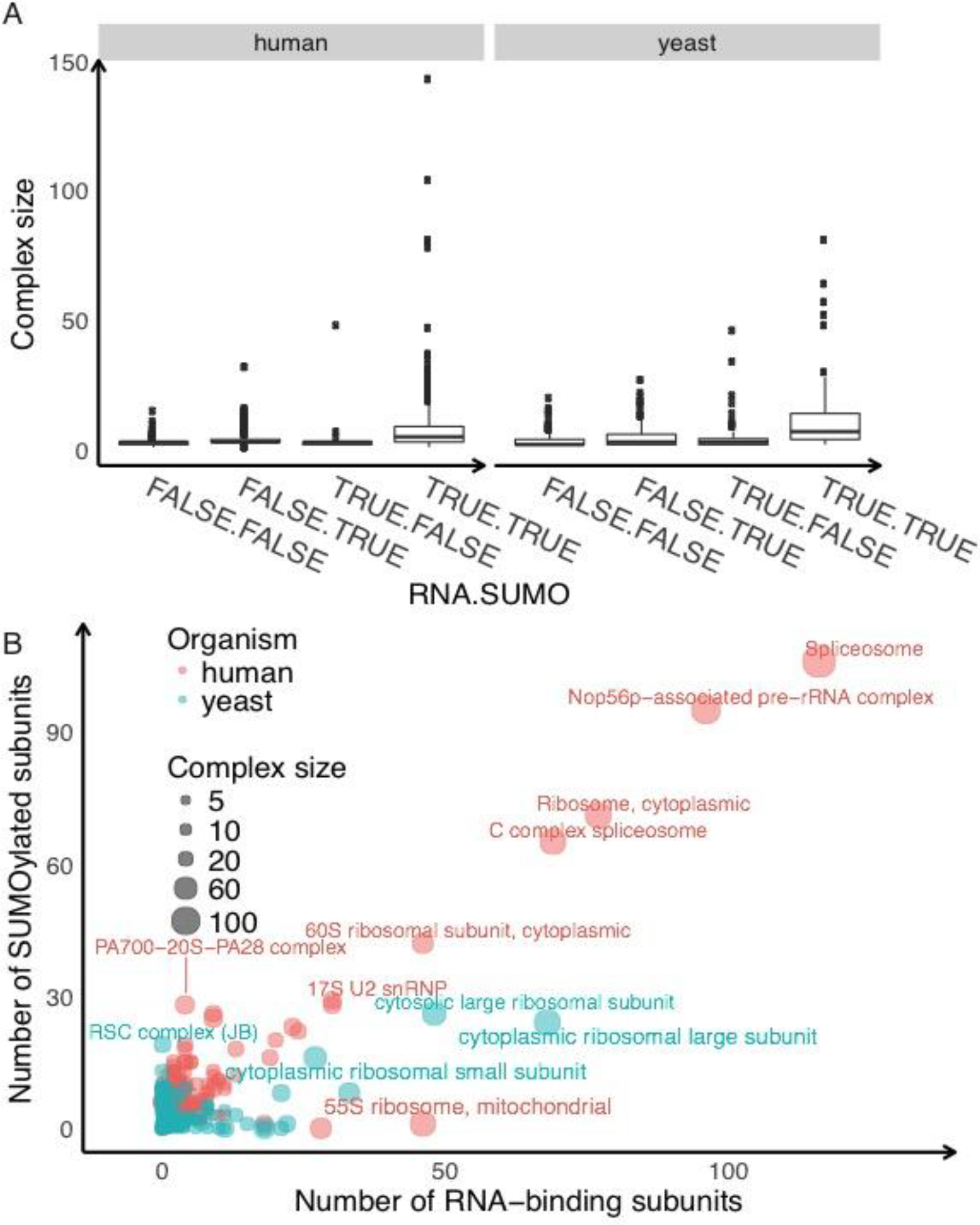
Correlation between protein SUMOylation, the total number of distinct subunits per complex, and the number subunits that bind RNA. Complex information was taken from the CORUM database ^64^. Dot size is proportional to the total number of distinct subunits of the complexes. Complexes that are both SUMOylated and contain many RNA-binding proteins are larger than other complexes. **A.** The columns are annotated as Is-RNA-binding-protein(TRUE/FALSE). Is-SUMOylated-protein(TRUE/FALSE). Protein complexes with RNA-binding members are more often SUMOylated than others. **B.** Relationship between SUMylation and RNA-binding proteins.

One of the outlying groups is high in SUMOylation but has few RNA binding proteins, represented by human PA700-20S-PA28 complex, proteasome, and yeast RSC (chromatin structure remodeling) complex (**Figure 2B**). They both consist of multiple different subunits which may legitimize the extensive SUMOylation observed across studies, while neither has RNA binding function. Another outlier contains many RNA-binding proteins but little SUMOylation. An example is the human 55S mitochondrial ribosome, contrasted by the 60S cytoplasmic ribosome, which is almost entirely SUMOylated. This may be the result of non-random subcellular localization of SUMOylation enzymes, or systematic bias in the high-throughput capture assays.

### Integrating diverse protein annotations substantially boosts prediction of SUMOylation (iSUMO)

The wide range of characteristics of SUMOylated proteins (see above) along with just a handful of SUMO E3 ligases highlight the need for tools that include more than sequence information to predict the SUMOylation state of proteins. To this end, we developed iSUMO, which employs a random forest model to make use of the intricate information in various protein annotations, ranging from basic molecular weight, SUMO-specific sequence motif, to biological functions, and connectivity in the protein-protein interaction network. This group of algorithms performs well with binary attributes which comprise much of the training set. In total, iSUMO integrates 638 and 293 attributes for human and yeast proteins, respectively (**Suppl. Table 1**). To show the benefit of incorporating protein annotations, we compared iSUMO to the same training method but with only sequence motif predictions from the well-established GPS-SUMO tool ^53^. We split the data into 65% training, 20% validation, and 15% testing subsets, and recorded training, 10-fold cross validation, validation performance metrics for model selection. Training methods varied with respect to termination condition, fixed number of trees, the maximum depth of trees, and whether we applied balancing of the training labels. We then chose the model with the highest area under the receiver-operator-characteristic (ROC) curve, and high F1, F2 values in validation and cross validation. In case of a tie, we chose the model with the shortest training time. To avoid overfitting, we also examined the model’s performance on test set. We accepted the model choice if it was close to training and validation metrics.

Overall, iSUMO showed a substantial improvement over predictions based on sequence alone (**Figure 3A**). For example, iSUMO’s average area underneath the ROC is 0.88 and 0.84 for human and yeast, respectively, compared to the sequence-based areas of 0.58 and 0.58. Further, at a 5% false positive rate (FPR), iSUMO’s true positive rate is about five-fold higher than that of the sequence-based predictions in human with 53% vs. 9%, respectively. This 5% FPR corresponds to an iSUMO score cutoff of 0.19 and 0.18 for human and yeast, respectively, which in turn predict 2,291 and 706 SUMOylated human and yeast proteins. The complete predictions, including for the extended human dataset, are available in the **Suppl. Table S6**.

**Figure 3.**
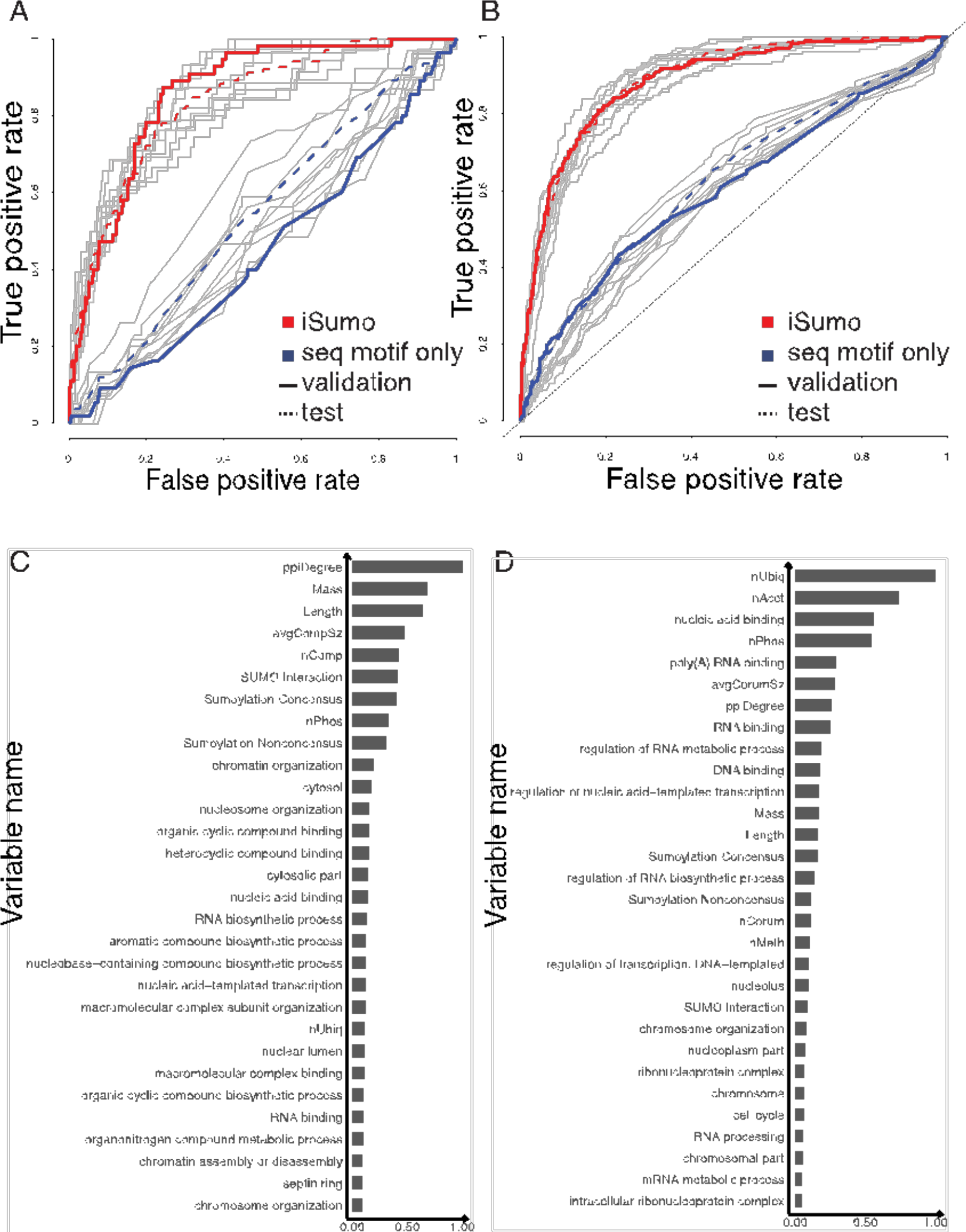
iSUMO predictions outperform sequence-based predictions. A., B. Receiver operator characteristics of iSUMO predictions trained on integrated sequence and annotation-based features (red) versus sequence-based features only (blue). Gray lines are the original 10-fold cross validation runs for different sets of non-overlapping, randomly chosen true negative entries. Balancing the number of positive and negative labels ensures learning quality and fair ROC evaluation. C., D. Frequencies of the most predictive attributes in human and yeast, measured as the number of occurrences in the different models. **Suppl. Table S4** lists all attributes used. ‘Depth’ marks the level in the decision tree and is displayed in different colors. The more frequent a feature is selected at low tree depth, the more predictive of SUMOylation it is.

### Predictive features of SUMOylation in yeast and human

Next, we analyzed the iSUMO models for attributes that are highly predictive (**Figure 3B**). From the random forest model training, we can estimate the relative importance of individual features by the frequency of being chosen at the split and the reduction of squared error it introduces. First, a protein’s connectivity in the PPI network (“ppiDegree”) ranks top in human model and fourth in yeast. Similarly, the complex size (“avgCompSz”) and number of complex memberships of a protein (“nComp”) rank fourth and fifth in the yeast model, respectively, and “avgCorumSz” and “nCorum” rank seventh and eleventh in the human model. These observations are consistent with the putative role of SUMOylation in supporting non-covalent protein-protein interactions and complex assembly and is likely common across eukaryotic organisms.

Second, the most important GO terms in both models are centered around DNA or RNA processing pathways, molecular function, or cellular localization (**Figure 3B**). As before, RNA binding and related functions are much more important in the human than in the yeast model. This finding suggests an increasing role of SUMOylation from transcriptional to post-transcriptional regulation in higher eukaryotes.

Third, unique to the human model, the numbers of ubiquitylation, acetylation, and phosphorylation sites in a protein are three highly predictive features, while they are much less important in the yeast model. This result is in line with numerous pieces of evidence on the extensive cross-talk between SUMOylation and other PTMs ^22, 54^. However, the pattern is less prevalent in yeast, perhaps because more data is available for human than yeast and the extent of post-translational modifications increases in higher organisms.

Fourth and last, basic molecular characteristics like length and weight as well as sequence motifs (consensus, non-consensus, and interaction) are far more dominant features in the yeast than in the human model (**Figure 3B**). This result gives a hint on how SUMOylation achieves it substrate specificity with a limited number of E3 ligases compared to, for example, kinases and ubiquitination enzymes. It is tempting to speculate that in the simpler yeast, SUMOylation only requires sequence specificity, but in the more complex mammalian system, SUMOylation requires cross-talk with other proteins and other protein modifications. Again, the trend might also be due to human data being complemented by a large number of additional information.

### Validation and application of iSUMO predictions

We used iSUMO with the high-quality reference proteomes that we created (**Method**) to provide a quantitative prediction of SUMOylation events in human and yeast that are highly enriched in functional SUMOylation events (**Suppl. Table 6a-c**). For high-confidence predictions, we restricted the analysis to well-annotated human genes and extracted predictions with the false positive rate (FPR) smaller than 0.05. The FPR was determined based on the model’s performance with a part of data unseen during training. At the 5% FPR threshold, iSUMO predicts 2,291 and 706 in human and yeast, respectively (**Suppl. Tables S6a, S6c**). Interestingly, the number is very similar to the combined number of experimentally determined SUMOylation events in the two organisms (2,270 and 527, respectively, **Suppl. Table S7**). Given a total size of the proteomes used for the study, ~19% and ~11% of the human and yeast proteome appears to be SUMOylated, respectively. This number is robust to the use of an extended dataset in human (**Suppl. Table 6b**).

At 5% FPR, iSUMO predicts SUMOylation of 467 human and 303 yeast proteins that were never observed in any of the 24 experimental datasets used for this study (**Suppl. Table S7**). To further validate these predictions, we searched other databases and literature for evidence of these SUMOylation of the top best predictions in human (**Table 3**). All ten proteins show some evidence for the prediction to be true. The yeast homolog of the ribosomal protein RRP1 is known to be SUMOylated ^55^, and for four proteins (ISY1, HIST2H2AB, TUBA3C, NSD1), other large-scale databases report SUMOylation events (Table 3). For other proteins, we found indirect evidence. For example, three of the proteins (SF3A3, ISY1, SNRPA) are part of RNA splicing, and the spliceosomal complex is known to depend on SUMOylation ^56^. Other proteins are part of chromatin reorganization and segregation during cell division (CHAF1B, ANAPC5, TUBA3C), which also involves many SUMOylation events ^57^. SUMOylation also plays a role in rRNA maturation and assembly of the 60S subunit ^9^, supporting the predicted SUMOylation of UTP15. Finally, plenty of literature supports SUMOylation of histones and histone methyltransferases ^58-59^, supporting the predicted modification of HIST2H2AB and NSD1. In sum, we argue that even these new predictions which had a high iSUMO score but were absent from the experimental training data, are highly enriched in true positives, i.e. functional modification events.

**Table 3.** Highest-scoring, newly predicted SUMOylation targets are likely true events. The table lists human proteins that have high iSUMO scores, but were not part of the training dataset. The annotation - through independent databases and literature search - shows that their SUMOylation is highly likely. The protein names are the primary common names as used by UniProtKB (http://www.ebi.ac.uk/reference_proteomes). The average iSUMO prediction scores range from 0 to 1, and is the higher, the higher the probability for SUMOylation. Function annotation is from GeneCards.org. * - SUMO site reported in PhosphoSitePlus ^65^; # - SUMO site reported in BioGrid ^66^. **Suppl. Table S6** shows the complete version of this table.

| Gene name | iSUMO score | Protein name | Protein function | Evidence for SUMO from literature |
| --- | --- | --- | --- | --- |
| **SF3A3 (SAP61)** | 0.82 | Splicing factor 3A subunit 3 | Subunit of the splicing factor SF3A (spliceosomal complex) required for 'A' complex assembly. | SUMOylation of spliceosomal protein complex is required for function <sup>56</sup>. |
| **RRP1 (NNP1, RRP1A)** | 0.82 | Ribosomal RNA processing protein 1 homolog A | Plays a critical role in the generation of 28S rRNA. | Yeast homolog is SUMOylated <sup>55</sup>. |
| **UTP15** | 0.71 | U3 small nucleolar RNA-associated protein 15 homolog | Ribosome biogenesis factor, involved in nucleolar processing of pre-18S ribosomal RNA. | SUMOylation, in particular via SENP3, plays a role in rRNA maturation and assembly of the 60S subunit <sup>9</sup>. |
| **ISY1 (KIAA1160)** | 0.71# | May play a role in pre-mRNA splicing | May play a role in pre-mRNA splicing. | SUMOylation of spliceosomal protein complex is required for function <sup>56</sup>. Sumoylation detected under heat stress <sup>67</sup>. |
| **CHAF1B (CAF1A, MPP7)** | 0.57 | Chromatin Assembly Factor 1 Subunit B | Complex that is thought to mediate chromatin assembly in DNA replication and DNA repair. | SUMO-2 target identified in screen. SUMOylation essential spindle organization, chromosome congression, and chromosome segregation <sup>57 50</sup>. |
| **SNRPA** | 0.55 | Small Nuclear Ribonucleoprotein Polypeptide A | Component of the spliceosomal U1 snRNP, which is essential for recognition of the pre-mRNA 5' splice-site and the subsequent assembly of the spliceosome. | SUMOylation of spliceosomal protein complex is required for function <sup>56</sup>. SUMOylation known for SNRPA1. |
| **HIST2H2AB** | 0.54#* | Histone H2A type 2-B | Core component of nucleosome. | Histones are heavily SUMOylated, in particular histone H3 and H4 <sup>58</sup>. |
| **ANAPC5 (APC5)** | 0.53 | Anaphase-promoting complex subunit 5 | Tetratricopeptide repeat-containing component of the anaphase promoting complex/cyclosome (APC/C), a large E3 ubiquitin ligase that controls cell cycle progression. | SUMOylation essential for spindle organization, chromosome congression, and chromosome segregation <sup>57</sup>. SUMO regulates mitosis and APC function in yeast <sup>68</sup>. |
| **TUBA3C (TUBA2)** | 0.53* | Tubulin alpha-3C/D chain | Major constituent of microtubules. | SUMOylation (especially also of tubulin) essential for spindle organization, chromosome congression, and chromosome segregation during cell division <sup>57 69 57</sup>. |
| **NSD1** | 0.51* | Histone-lysine N-methyltransferase | Methylates 'Lys-36' of histone H3 and 'Lys-20' of histone H4 (in vitro). | SUMO regulates some methyltransferases <sup>59</sup>. |

## Discussion

A major question arising from these comparative studies concerns the total number of SUMOylated proteins. Here, we present iSUMO, an integrated SUMOylation prediction framework that outperforms methods based on sequence features alone (**Figure 3**). We apply iSUMO to both human and yeast proteins, predicting 2,291(19%) and 706 (11%) of SUMOylated proteins, respectively. Encouragingly, this number is close to a recent estimate by Hendriks et al. which, combining several experimental studies, predicted a total set of ~3,600 (15%) of SUMOylated human proteins ^60^.

However, a remaining challenge in these analyses is the small overlap between the individual large-scale studies, suggesting that each experiment produces many false positive identifications (**Figure 1**). Alternatively, our current methods might be unable to capture the complete SUMO space and only detect small subsets. Therefore, the next goal should be to identify the true positive SUMOylation events amongst the many detected ones. iSUMO offers a solution to this problem by integrating sequence and protein functional features to learn computationally the properties of SUMOylated proteins. It outperforms prediction tools that are solely based on sequence features and produces a list of high-confidence predictions. In total, 1,824 (15%) and 403 (6%) proteins in human and yeast, respectively, are positive iSUMO predictions *and* occur in the experimental datasets (**Suppl. Table S7**) – these proteins are most likely to be truly functional.

In addition, our study also highlighted protein characteristics strongly connected to SUMOylation. For example, SUMOylated proteins often bind nucleic acids (e.g. RNA or DNA) and are part of large complexes (**Table 2**)^22^. Specifically, there is strong correlation between SUMOylation, the size of a complex, and the number of subunits that are RNA-binding (**Figure 2**). Overall, two fifths (654 of 1,536) of the human RNA-binding proteins are SUMOylated, while this is the case for less than 10% of the total human proteome -- suggesting that SUMOylation might play a role specifically in mediating protein-RNA binding, beyond its known function as a facilitator of protein-protein interactions. Whether SUMOylation modifies the structure of the RNA-binding protein, or affects its surface charge to enable the interaction with the nucleic acid remains subject to future studies.

It is tempting to speculate on the reasons for the prevalence for SUMOylation amongst RNA-binding proteins. Perhaps, with the expansion of RNA-based regulatory pathways in mammals compared to yeast, the well-established, extensive role of SUMOylation of DNA-binding proteins was simply transferred. Alternatively, SUMOylation might be essential for the correct assembly of large complexes, which are very often involved in RNA-related processes, and the prevalence of SUMOylation for RNA-binding proteins might be a side-effect of its role in complexes. A third intriguing hypothesis arises from two observations: SUMO is one of the most soluble of all known proteins ^61^ and RNA-binding proteins are major components of RNA-protein granules whose aggregation forms the molecular bases of many neurodegenerative disorders^62^. Therefore, SUMOylation may act to prevent such aggregation in these densely packed cellular structures – a hypothesis supported by some experimental work ^63^.

## Acknowledgements

C.V. acknowledges funding by the NIH (Ro1 GM113237), the NSF EAGER grant, the DOD (Hypothesis Testing Award PC121532), and the NYU Whitehead Foundation. X.Y. acknowledges funding by the Biology Department at New York University (Master’s Research Grant).

## Supporting Materials

Supplementary Tables in Excel file.

S1a. Human proteome-wide SUMOylation data

S1b. Yeast proteome-wide SUMOylation data

S2a. Enriched GO categories, human

S2b. Enriched GO categories, yeast

S3. Protein complexes, human

S4. Features used in modeling

S5. Model selection, human and yeast

S6a. Final prediction, human - high-confidence dataset

S6b. Final prediction, human - extended dataset

S6c. Final prediction, yeast

S7. Numbers

